# Landscaping the behavioural ecology of primate stone tool use

**DOI:** 10.1101/2021.08.24.457369

**Authors:** Katarina Almeida-Warren, Tetsuro Matsuzawa, Susana Carvalho

## Abstract

Ecology is fundamental to the development, transmission, and perpetuity of primate technology. Previous studies on tool site selection have addressed the relevance of targeted resources and raw materials for tools, but few have considered the broader foraging landscape. In this first landscape-scale study of the ecological contexts of wild chimpanzee (*Pan troglodytes verus*) tool-use, we investigate the conditions required for nut-cracking to occur and persist over time at discrete locations in Bossou (Guinea). We examine this at three levels: selection, frequency of use, and inactivity. We find that, further to the presence of a nut tree and availability of raw materials, abundance of food-providing trees as well as proximity to nest sites were significant predictors of nut-cracking occurrence. This suggests that the spatial distribution of nut-cracking sites is mediated by the broader behavioural landscape and is influenced by non-extractive foraging of predictable resources, as well as non-foraging activities. Additionally, the number of functional tools was greater at sites with higher frequency of nut-cracking and was negatively correlated with site inactivity. Our findings indicate that the technological landscape of the Bossou chimpanzees shares affinities with the ‘favoured places’ model of hominin site formation and provides new insights for reconstructing ancient patterns of landscape use.

**Résumé:** L’écologie est fondamentale pour le développement, la transmission et la pérennité de la technologie des primates. Des études antérieures ont identifié la disponibilité des ressources cibles ainsi que les matières premières pour les outils comme des facteurs influents dans la sélection des emplacements pour les activités technologiques. Cependant, il y a peu d’études qui abordent cette recherche à l’échelle du paysage et du comportement fourrager. Dans cette première étude paysagère sur l’utilisation d’outils par le chimpanzé sauvage (*Pan troglodytes verus*), nous recherchons les conditions écologiques qui influencent la sélection, l’utilisation et l’inactivité des emplacements utilisés pour le cassage des noix en Bossou, Guinée. Nos résultats montrent qu’en plus de la présence d’un noyer et de la disponibilité des matières premières, l’abondance d’arbres nourriciers ainsi que la proximité des sites de nidification étaient des prédicteurs significatifs de l’occurrence du cassage des noix. Cela suggère que la distribution spatiale des sites de cassage de noix est influencée par le paysage comportemental et est influencée par le fourrage non-extractive de ressources prévisibles, ainsi que par des activités non-fourragers. Nos résultats indiquent que le paysage technologique des chimpanzés de Bossou partage des affinités avec le modèle des « lieux favoris » de la formation des sites hominidés et fournit de nouvelles perspectives pour reconstruire les modes d’utilisation du paysage anciens.

## Introduction

Ecology plays an important role in shaping non-human primate behaviour from foraging strategies, to ranging patterns, and sociality (Robbins and Hohmann, 2006; Strier, 2011). Tool-use, particularly for extractive foraging, is no exception. Recent studies have highlighted that ecology is key in determining whether tool-use emerges in a population, how it manifests itself, and how it is maintained once it is established (S. Carvalho et al., 2007, 2011; Grund et al., 2019; Koops et al., 2013, 2014).

Stone tool-use is often recurrent in spatially discrete locations, frequently involves the reuse of tools, and leaves a recognisable archaeological footprint that can be traced back thousands of years (Falótico et al., 2019; Mercader et al., 2007). However, little is known about the ecological factors influencing selection and repeated use of specific locations for these activities and how they fit within the broader foraging landscape. Lithic-based foraging technology has been recorded in several wild, non-human primate species including chimpanzees (*Pan troglodytes ssp.*; Whiten *et al*., 1999), bearded capuchin monkeys (*Sapajus libidinosus;* Ottoni and Izar, 2008), Burmese long-tailed macaques (*Macaca fascicularis*; Gumert, Kluck and Malaivijitnond, 2009), and, most recently, in white-faced capuchins (*Cebus capuchinus*; Barrett *et al*., 2018).

Chimpanzees are of particular interest because they are our closest living relatives (Langergraber et al., 2012), and they present the largest, most diverse and ecologically adaptable technological repertoire compared to any other non-human species, reflecting a level of cognitive flexibility akin to the earliest hominin toolmakers (S. Carvalho et al., 2013; Pascual-Garrido and Almeida-Warren, 2021; Rolian and Carvalho, 2017). Chimpanzee nut-cracking assemblages have been found to have close similarities to the low-density assemblages characteristic of the early hominin record (S. Carvalho et al., 2008; S. Carvalho and McGrew, 2012). Thus, understanding how patterns of nut-cracking behaviour accumulate across the landscape can provide valuable insights into the formation and spatial distribution of early hominin assemblages and allow the modelling of ancient landscape use and resource exploitation.

Previous research on chimpanzee nut-cracking has established that the spatial availability of nut trees and raw materials for tools influences site location and reuse, as well as frequency and distance of tool transport (S. Carvalho et al., 2007, 2011). Nevertheless, nut-cracking assemblages are yet to be explored within the context of the broader ecological and foraging landscape. This requires the study of nut-cracking sites not only in relation to direct ecological correlates such as access to raw materials and nuts, but also in relation to ecological requirements of other daily activities critical to survival such as food, water, and shelter.

Chimpanzees living in forested environments spend approximately 50% of their waking hours foraging and travelling between feeding locations (Pruetz and Bertolani, 2009). Recent studies have shown that chimpanzee ranging patterns are dynamic and are influenced by the spatial distribution and seasonality of food (Trapanese et al., 2019), which may also determine where non-foraging activities, such as nesting, take place (Basabose and Yamagiwa, 2002; Hernandez-Aguilar, 2009; Janmaat et al., 2014). Their diet mainly consists of fruit (Morgan and Sanz, 2006), but high-energy foods such as insects, nuts and honey acquired through tool-assisted foraging are also important staples or nutritional supplements for many chimpanzee populations (Sanz and Morgan, 2013). Yet, little is known about how extractive foraging interacts with other feeding activities and the broader behavioural landscape. This study explores this question for the first time in the context of chimpanzee nut-cracking using stone tools.

Water is essential to life (Popkin et al., 2010). For non-human primate species living in extremely arid conditions, such as savannah-dwelling chimpanzees and baboons, water is a critical resource that constrains movement patterns and landscape use (Barton et al., 1992; Pruetz and Herzog, 2017; Wessling et al., 2018). It has also featured in many discussions surrounding early hominin evolution and behaviour (Joordens et al., 2019), and has been spatially linked with early stone tool sites (Rogers et al., 1994). Similarly, the location of chimpanzee nut-cracking sites has been suggested to coincide with the proximity to hydrological features, such as streams and rivers (S. Carvalho et al., 2007), possibly because they are the most likely sources of eroded lithic raw materials for tools (S. Carvalho, 2011), although this remains to be empirically tested.

Chimpanzees habitually make a sleeping nest at the end of every day, and sometimes make day nests for resting (Koops et al., 2012). They have been an important focus of research since Sept (1992) recognised that they form clusters of debris akin to early hominin assemblages, and saw their potential for understanding patterns of early hominin landscape use and the origins of human shelter (Mcgrew, 2021). Nesting locations have subsequently been linked with a range of ecological parameters such as tree species and tree architecture, as well as surrounding topography and vegetation types (Badji et al., 2018; J. S. Carvalho et al., 2015; Hernandez-Aguilar, 2009; Hernandez-Aguilar et al., 2013; Koops et al., 2012; Ndiaye et al., 2018; Stewart et al., 2011). Other studies have found additional links between nest sites and proximity to areas of high fruit availability (Basabose and Yamagiwa, 2002; Furuichi and Hashimoto, 2004; Goodall, 1962; Janmaat et al., 2014). While the exact conditions determining nest site suitability appears to be population-specific rather than universal, these findings demonstrate that nesting activities shape chimpanzee landscape use, and, in turn, are shaped by resource distribution and the local environment. Nevertheless, little is known about how nesting relates to other activities such as spatially discreet forms of tool use (e.g., nut-cracking and termite-fishing).

This is the first landscape-scale investigation of the ecological drivers of chimpanzee tool-use, where we examine the conditions required for nut-cracking to occur and persist over time in discrete locations at the long-term field site of Bossou (Guinea). This is divided into three points of enquiry: 1) tool site selection; 2) tool site use; 3) tool site inactivity.

*Tool site selection* explores the ecological conditions that determine where nut-cracking sites are established within the chimpanzee home-range. This is critical to understanding how technological activities occur within the broader behavioural landscape, and how these activities produce locally discrete assemblages that can remain archaeologically identifiable for thousands of years (e.g. Falótico et al., 2019; Mercader et al., 2007). There is a growing body of research on habitat selection for daily activities such as foraging, travelling, socializing and sleeping, particularly within the context of anthropogenic landscapes and the implications for conservation (Bryson-Morrison et al., 2017; K. B. Potts et al., 2016). However, regarding technological activities, although we are beginning to learn more about raw material selection for tools (S. Carvalho et al., 2008; Pascual-Garrido and Almeida-Warren, 2021), the selection of locations for tool use remains unexplored.

*Tool site use* focuses on the ecological factors that influence the frequency with which established sites attract nut-cracking activity. How often a site is used can serve as a proxy for inferring preference in relation to other sites. From an archaeological perspective, generally the more a location is used for debris-generating activities, the larger and more conspicuous an archaeological signature becomes. This has important implications for understanding the spatial clustering of activities and material evidence over time and can offer many insights into the formation and perpetuity of early hominin archaeological assemblages (S. Carvalho and Almeida-Warren, 2019; McGrew, 2010). Ethoarchaeological studies of chimpanzee nests have found that sleeping sites are frequently revisited and the nests themselves may be reused (Hernandez-Aguilar, 2009; Sept, 1992; Stewart et al., 2011). However, comparable literature on the use of tool sites is scarce, except for the reuse of stone tools (S. Carvalho et al., 2009), and sources of perishable raw materials (Pascual-Garrido, 2018; Pascual-Garrido and Almeida-Warren, 2021).

*Tool site inactivity* investigates the ecological conditions that may cause the cessation of nut-cracking activity at an established tool site. Chimpanzees live in dynamic landscapes shaped by environmental change. Whether natural or anthropogenic, these shifts often result in changes in resource distributions that may in turn lead to the development of new behavioural adaptations and ranging patterns (e.g. Kalan et al., 2020). New opportunities may arise (e.g., foraging food from crops: Hockings et al., 2015, 2012; nesting in novel plant species: McCarthy et al., 2017), while formerly habitual activities may shift to new locations or become obsolete (Gruber et al., 2012; Kühl et al., 2019). Environmental changes have been identified as important drivers of our own evolutionary history (Bobe et al., 2002; Bobe and Carvalho, 2019; Joordens et al., 2019; R. Potts, 1998; R. Potts et al., 2020; Reed, 1997), but few studies have addressed empirically how these changes may have affected patterns of landscape use and the distribution of early hominin tool sites (e.g. Rogers et al., 1994). Thus, investigating the conditions that influence the cessation of activity at chimpanzee tool sites, can provide important clues as to the factors that lead to their temporary or long-term abandonment.

Throughout these steps we assess the effect of ecological parameters that have been found to correlate with nut-cracking activities (nut availability; abundance of raw materials; distance to water) as well as variables that encapsulate two key aspects of chimpanzee activity patterns: non-extractive foraging (abundance of food resources: wild food trees; wild fruit trees; THV - terrestrial herbaceous vegetation) and sleeping (distance to nesting site).

## Methods

### Study site and subjects

Bossou (7° 39’ N, 8° 30’ W) is located in the southeast of the Republic of Guinea (West Africa), 6 km from the foothills of Mount Nimba Strict Nature Reserve (Figure 1) (Humle, 2011a; Yamakoshi and Sugiyama, 1995). The chimpanzee community has been studied continuously in both natural (since 1976) and experimental (outdoor laboratory since 1988) settings (Tetsuro Matsuzawa et al., 2011; Sugiyama and Koman, 1979). Between 1976 and 2003 the population size ranged from 18 to 23 individuals (Sugiyama, 2004), but has since declined largely due to a catastrophic flu-like epidemic from which it never recovered (Humle, 2011b; Sugiyama and Fujita, 2011). At the time of this study, the population consisted of seven individuals. The Bossou chimpanzees habitually crack and consume oil-palm nuts (*Elaeis guineensis*) and are currently the only population known to use portable stones as both hammers and anvils (Carvalho, Matsuzawa and McGrew, 2013; but see Ohashi, 2015 for recent discoveries in Liberia). Bossou has two seasons – a short dry season lasting from November to February, and a long rainy season extending from March to October (Humle, 2011a; Yamakoshi, 1998). Nut-cracking occurs year-round, but is most prevalent during peak wet season (June – August) and at the start of the dry season (November – December) when fruit is less abundant (Yamakoshi, 1998).

**Figure 1.**
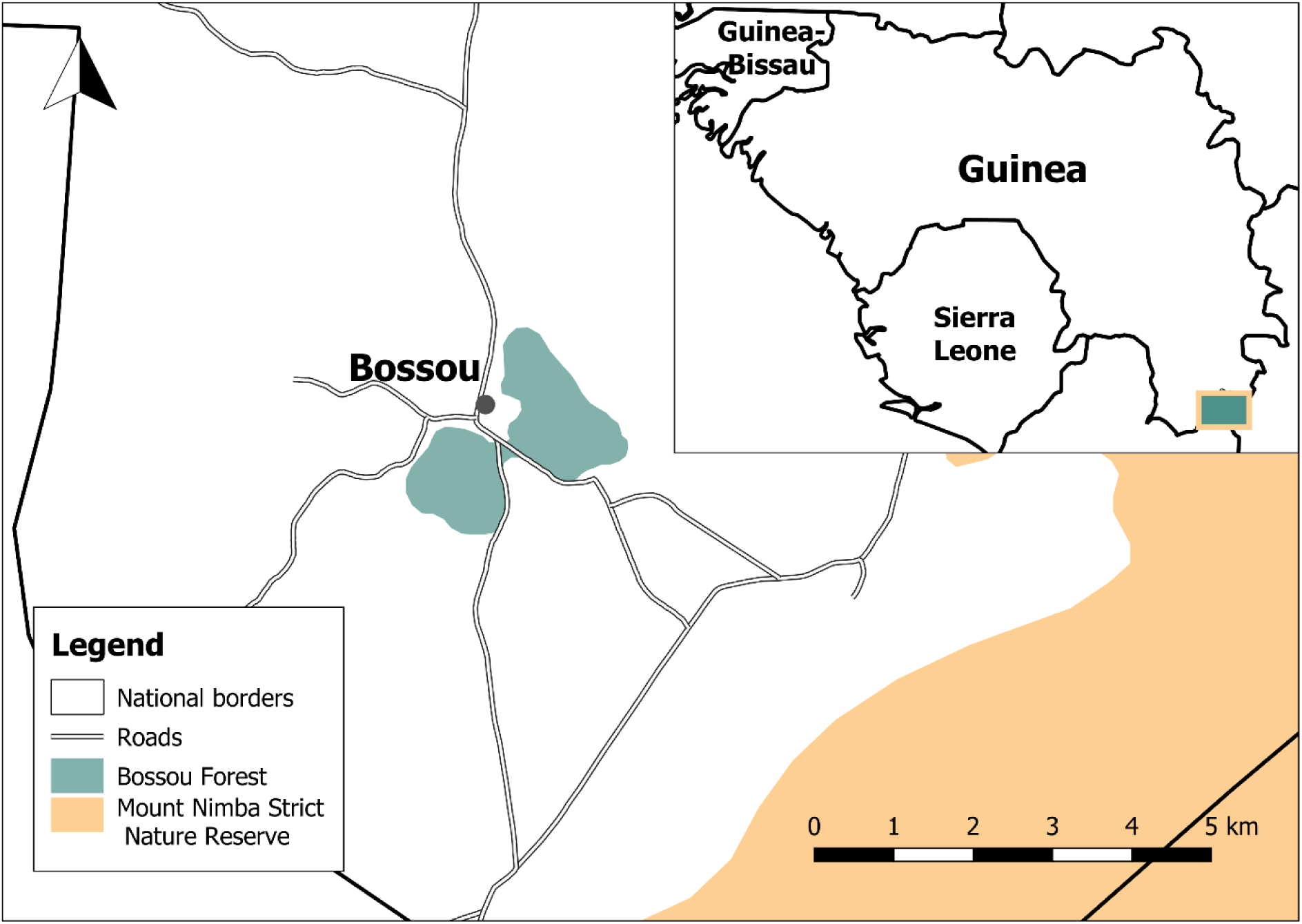
Map of the Bossou Forest and surrounding area. Mount Nimba Strict Nature Reserve shapefile adapted from the WDPA database (UNEP-WCMC and IUCN, 2019)

The Bossou forest has an estimated area of 16 km^2^ and is intersected by roads (Figure 2; Hockings, Anderson and Matsuzawa, 2006). Within this, the chimpanzees range a core area of approximately 7 km^2^ (Hockings et al., 2006). The habitat is comprised of a composite of primary, secondary and riverine forests, savanna and cultivated fields (Hockings et al., 2012). The northern slope of Mont Gban, in the Eastern part of the forest, is considered a sacred area – forêt sacrée – by the local *Manon* culture, and access is forbidden to outsiders. Out of respect to this tradition, no research was conducted in this area.

**Figure 2.**
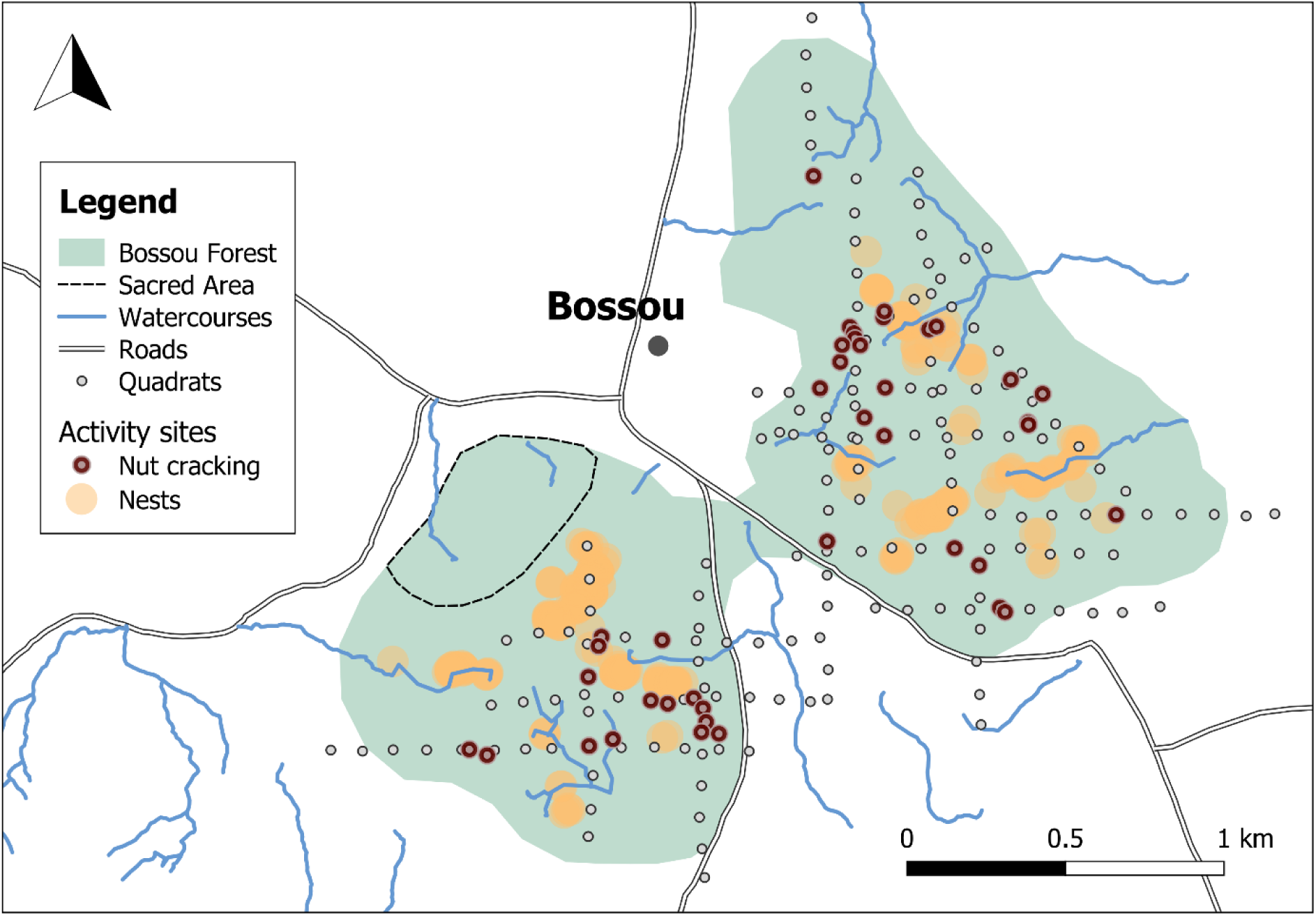
Map of the study area and surrounding area, highlighting the locations of recorded nut-cracking sites, quadrats, nests, and watercourses.

### Data collection

Data were collected over two field trips: 14DEC17-01MAY18 and 28OCT18-13DEC18, encompassing 160 days of fieldwork. We employed a mixed-method approach that combined direct behavioural observation through active group follows of the chimpanzee population, with archaeological documentation of nut-cracking sites (indirect behavioural observations), and ecological research using the transect and quadrat method. At an initial stage, we targeted nut-cracking sites that had previously been documented by SC in 2006 and 2008-09. Further nut-cracking sites were discovered during surveys and group follows throughout both research seasons. For each nut-cracking site we established 1-km transects intersecting the site datum at 500 metres. Nut-cracking sites within 100 metres of a pre-established transect were either assigned to that transect or became the mid-point of a new perpendicular transect, to ensure even forest coverage. All transects were oriented N-S or E-W, except for two that were oriented NE-SW and NW-SE due to access difficulties (Figure 2). 5-metre radius survey quadrats were established at every 100-metres along the transects starting from the midpoint where the nut-cracking site was located. At each quadrat nut-cracking specific and general ecological and vegetation data were collected (further details below).

We employed a fully digital method of data collection. Quadrat datums and all data entries (food-providing vegetation, tools, raw materials) were georeferenced using an Arrow Gold GNSS receiver (µ_HRMS_ = 2 metres; Almeida-Warren et al., 2021; EOS Positioning Systems Inc., 2017). Coordinates were instantly downloaded via Bluetooth to the GeoGrafi-M application (MGISS, 2019) on an android device where further data could be entered through custom made forms.

#### Oil palms

For each oil palm encountered during quadrat surveys we documented diameter at breast height (DBH), and number of fruit bunches (total and ripe). For 25 of the nut trees associated to nut-cracking sites we also collected information on nut availability and new traces of nut-cracking on a weekly basis during the first field season (22JAN18-03MAY18) and once at the beginning and end of the second season (weeks of 29OCT18; 10DEC18). Additional data was collected by Henry Camara during the weeks of 30SEP19, 27APR20, and 25MAY20. As per Koops *et al*. (2013), we scored presence of edible nuts on the ground within a 2-metre radius of the nut tree: (0) nuts absent; (1) 1-50 nuts; (2) 51-100 nuts; (3) > 100 nuts. With aid from field guides, nut suitability was determined by checking a sample of randomly collected nuts for whether the nuts contained an edible kernel or were rotten (following Koops *et al*., 2013). Nuts were not opened so as not to affect future availability, but the local people also crack oil palm nuts and are able to identify whether or not they are edible (Humle and Matsuzawa, 2004).

#### Tools and raw materials

All lithic material was recorded for size, raw material type, and portability (whether loose or imbedded in the ground). Adapted from Koops *et al*. (2013), size was scored into six categories: (1) 1-2 cm; (2) 3-5 cm; (3) 6-10 cm; (4) 11-20 cm; (5) 21-30 cm; (6) >30 cm. Tools and bi-products of nut-cracking were defined as stones that showed at least one of the following: a) traces of wear from nut-cracking; b) nutshell remains on or around them; c) could be refitted with another stone with evidence of a) or b). For this study, the variable *Tools* included all lithic materials used for nut-cracking excluding fragments that no longer or could no longer be used for nut-cracking.

The collective tool assemblage was also scored for status of nut-cracking activity: (Active) New signs of nut-cracking activity were recorded during the fieldwork period. Nut powder or cracked nut kernels were visible on top of or around tools, and there was at least one hammer and anvil pair with impact points that had not rusted over; (Inactive) There were no signs of recent nut-cracking activity during the entire fieldwork period. Cracked nut kernels were either absent or present but showed clear signs of decay. Iron oxide or moss developing on tool impact points.

#### Vegetation

We recorded all wild non-THV plants with a DBH > 2cm of species known to be consumed by the chimpanzees of Bossou. We then cross-referenced the recorded species with the current list of chimpanzee food resources to identify those which were sources of fruit – the preferred food-type of chimpanzees. To distinguish permanent food sources from ephemeral food sources, THV was documented separately. Domesticated or crop species were not included in this study.

#### Nests

Because few nests were documented at the quadrat level or along transects, additional forest-wide surveys were conducted through random walks and strategic walks targeting areas that were known nesting locations. When a nest was encountered, elevation and direction of travel were maintained along topographical contours and all nests within 50-metres of the nest and either side of the projected route were documented until no further nests were visible over a 50-metre stretch (modified from Hernandez-Aguilar, 2009). For data analysis, nests were divided into spatial clusters. A cluster was defined as a 50-metre radius area with a minimum of 21 documented nests, representing a minimum of three sleep-events for the collective chimpanzee population (N = 7) as a proxy for a habitual nesting location. This was achieved on QGIS (Version 2.18.4), by creating a 50-metre buffer zone around each nest and using the ‘count points in polygon’ function to identify zones with a minimum of 21 nests. The *distance to nearest nest cluster* variable used in the following analyses was computed with the ‘distance to nearest hub’ function and was defined as the distance from a quadrat datum to the centre of the nearest nest cluster.

#### Watercourses

Watercourse data was collected in 2008/2009 by SC. With the help of field guides, streams and rivers were traced on foot and recorded using the track feature on a Garmin hand-held GPS device. The *distance to nearest river* variable used in the following analyses was computed in QGIS and was defined as the distance between a quadrat datum to the nearest point along the watercourse polylines.

### Data analysis

#### Tool site selection

Initial inspection of the data revealed that no nut-cracking occurred in quadrats where nut trees were absent. This is consistent with previous literature describing that nut-cracking occurs in close proximity to a nut tree (S. Carvalho et al., 2008). Additionally, there was very little variation in the number of oil palms in each quadrat with only 15% of oil palm quadrats documented with more than one oil palm. For these reasons, we restricted data analysis to quadrats where an oil palm was present and did not include the number of oil palms as a predictor, as this would mask the potential effect of the other variables of interest. The final dataset had a total of 82 quadrats, 40 of which had traces of nut-cracking activity. We used a binomial generalized linear model (GLM; Zuur *et al*., 2009a) with a logit link function to investigate the effect of five main predictors: raw materials, wild food trees, wild food THV, distance to nearest nest cluster, distance to nearest river, on the presence (1) versus absence (0) of a tool site in a given quadrat. We also analysed three sub-models to determine whether more restricted variables yielded a better model fit (Appendix, Table A2). The first sub-model replaced raw materials with a subset of raw materials of size class corresponding to the three most common tool size classes (95% of tools. Size class: 3, 4, 5; Appendix, Table A1). The second sub-model replaced wild food-providing trees with a subset formed only of fruit-providing trees. The third sub-model included raw materials of size class 3-5 and fruit-providing trees. Akaike’s Information Criterion for small sample sizes (AICc) was used to compare models (Burnham and Anderson, 2004), whereby the model with the lowest AICc was chosen as the final model.

#### Tool site use

We investigated whether the hypothesized ecological variables (i.e., nut availability, raw materials, food trees, distance to nearest nest cluster, and distance to nearest river) influenced the frequency a nut-cracking site was used. From a total of 361 monitoring observations, only 35 cases of recent nut-cracking events were identified for 17 out of the 25 monitored nut-cracking sites, where frequency of recent activity ranged between 1 and 4. Because of single (N = 1) sample sizes for 2 and 4 events, frequency of activity was recoded as “Low” (≤ 2 events; N = 10) and “High” (> 2 events; N = 7). The small sample size (N = 17) was deemed too small to justify a GL(M)M, therefore we only discuss descriptive statistics for this question using two-sample t-tests (or Man-Whitney U tests when assumptions of normality were not met).

#### Tool site inactivity

We used a binomial GLM with ‘logit’ link to investigate the effect of mean nut availability, raw materials, and food trees, on tool-site inactivity. The response variable included nut-cracking sites that were classified as active (response = 0; N = 24) with those classified as inactive (response = 1; N = 16). The final dataset included 40 tool sites. Akin the model for tool site selection, we also investigated four sub-models with raw materials of size class 3-5, tools, and fruit tree subsets (Appendix, Table A6). The model with the lowest AICc was then chosen as the final model.

#### General considerations

All analyses were processed in R Studio (version 1.1.383; R Studio Team, 2016), using R (version 4.1.0; R Core Team, 2021). Data exploration for each GLM, following the protocol described in Zuur, Ieno & Elphick (2010), did not raise any concerns. Collinearity among the explanatory variables was assessed by calculating the Variance inflation factors (VIF) using the function ‘vif’ of the car package (Jon Fox and Weisberg, 2011). None of the models indicated any multicollinearity issues (Maximum VIF = 1.39, Quinn and Keough, 2002). To assess the significance of the full models and sub-models, we ran likelihood ratio tests (LRT) using the ‘anova’ function which compared each model to a corresponding null model from which all fixed effects were excluded (Dobson, 2002). We tested the significance of main effects for each model by systematically dropping them one at a time and comparing the resulting model with the full model using the ‘drop1’ function (Dobson, 2002). P-values for the individual effects were based on the LRT results from the ‘drop1’ function. The AICc for model selection was calculated using the ‘MuMIn’ package (Bartón, 2020). Model assumptions were verified by plotting residuals versus fitted values and versus each covariate in the model (Zuur and Ieno, 2016). Influential observations were assessed by calculating and plotting the Cook’s distance (Smith and Warren, 2019); all values were under the recommended threshold of 1, suggesting no evidence of influential points (John Fox, 2002; Smith and Warren, 2019).

For tool site use, comparisons between low and high frequency of nut-cracking activity were computed for each of the variables of interest using unpaired two-sample t-tests or the non-parametric equivalent, Wilcoxon rank-sum test (i.e., Mann-Whitney U test). Normality assumptions were assessed using the Shapiro-Wilk test. Threshold for statistical significance was set to *p* ≤ 0.05.

### Data availability

The datasets generated during and/or analysed during the current study will be available from the corresponding author on reasonable request. Source code will be made available at https://github.com/katarinawarren/bossou-chimps-analysis [to be replaced with DOI].

### Ethical statement

All tool site and ecological data were collected when chimpanzees were absent from the survey locations. Efforts to ensure minimal disturbance of nut-cracking sites included: keeping all tools in their original locations; not removing or cracking nuts; collecting stone samples from existing tool fragments whenever possible. Research was conducted in accordance with all the research requirements of Guinea, and the ethical protocols set out by The University of Oxford, the Kyoto University Primate Research Institute, and the Institut de Recherche Environmentale de Bossou (IREB).

## Results

### Tool site selection

The sub-model where raw materials were replaced by a subset of size class 3-5 was the best fitted model according to the AICc (Appendix, Table A3), and had a clear effect on the probability of a nut-cracking site occurring in a location where at least one oil palm was present (full-null model comparison, LRT: *df* = 5, *deviance* = 56.52, *p* < 0.001). Raw materials had a significant positive effect on tool site prediction, as did food trees, while distance to nest cluster had a significant negative effect (Table 1; Figure 3). All other fixed effects were non-significant (Table 1). The sub-model replacing wild food trees with the fruit trees subset yielded the worst model fit in which fruit trees were not a significant predictor (Appendix, Table A3, Table A4).

**Figure 3.**
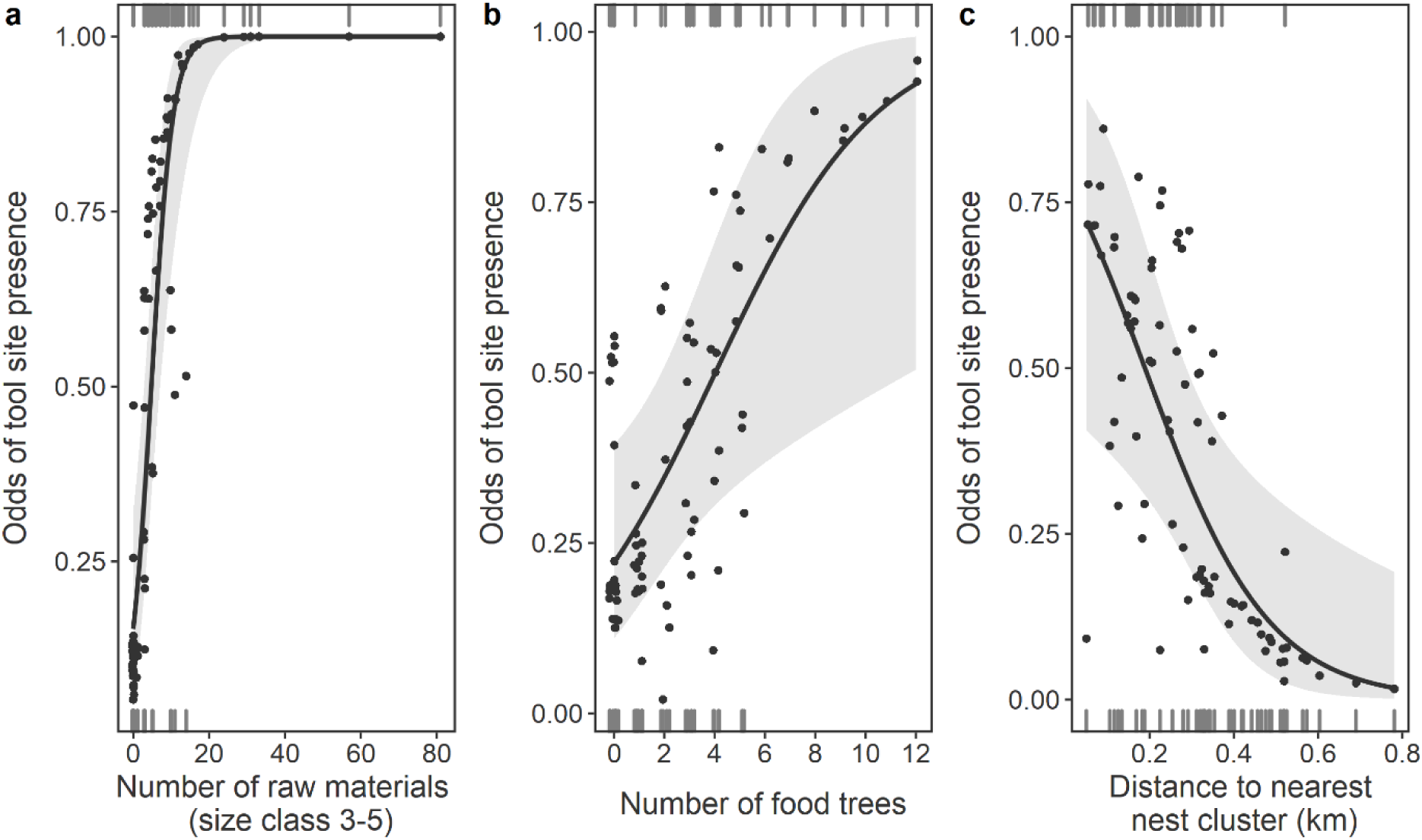
Probability of tool site presence in response to: a) Raw materials of size class 3 – 5; b) Trees that are sourced by chimpanzees for food; c) Distance to the nearest nest cluster.

**Table 1.** Results of the final GLM investigating potential predictors influencing tool site selection

| Term | Estimate | SE | $\chi^2$ | $p^a$ |
| --- | --- | --- | --- | --- |
| Intercept | -1.139 | 0.472 |  | NI<sup>b</sup> |
| Raw materials (size class 3-5) | 0.358 | 1.298 | 21.929 | <0.001 |
| Wild food trees | 0.313 | 0.407 | 5.758 | 0.016 |
| THV | -0.103 | 0.383 | 0.392 | 0.531 |
| Distance to nearest nest cluster | -0.007 | 0.485 | 6.109 | 0.013 |
| Distance to nearest river | 0.005 | 0.411 | 1.397 | 0.237 |

<sup>a</sup> Results from the likelihood ratio test using the 'drop1' function.
<sup>b</sup> Not indicated because it has a limited interpretation.

### Tool site use

Over a total period of 15 weeks, only 33 cases of nut-cracking were recorded for 17 out of 25 monitored tool sites. 10 of the 17 sites recorded one or two nut-cracking events (low frequency), with the remaining seven showing recent traces between three and four times during the monitoring period (high frequency). In general, mean nut availability was significantly higher at nut-cracking sites that registered a higher frequency of nut-cracking activity (T-test: *p* = 0.03; Figure 4; Table A5). Furthermore, distance to nearest nest cluster revealed a negative trend, whereby high frequency sites tended to be nearer to nesting locations (Wilcoxon rank-sum test: *p* = 0.07). For all other variables of interest (raw materials, mean fruit bunches, wild food trees and distance to nearest river) there were no significant differences between the groups (*p* > 0.15; Figure 4; Table A5).

**Figure 4.**
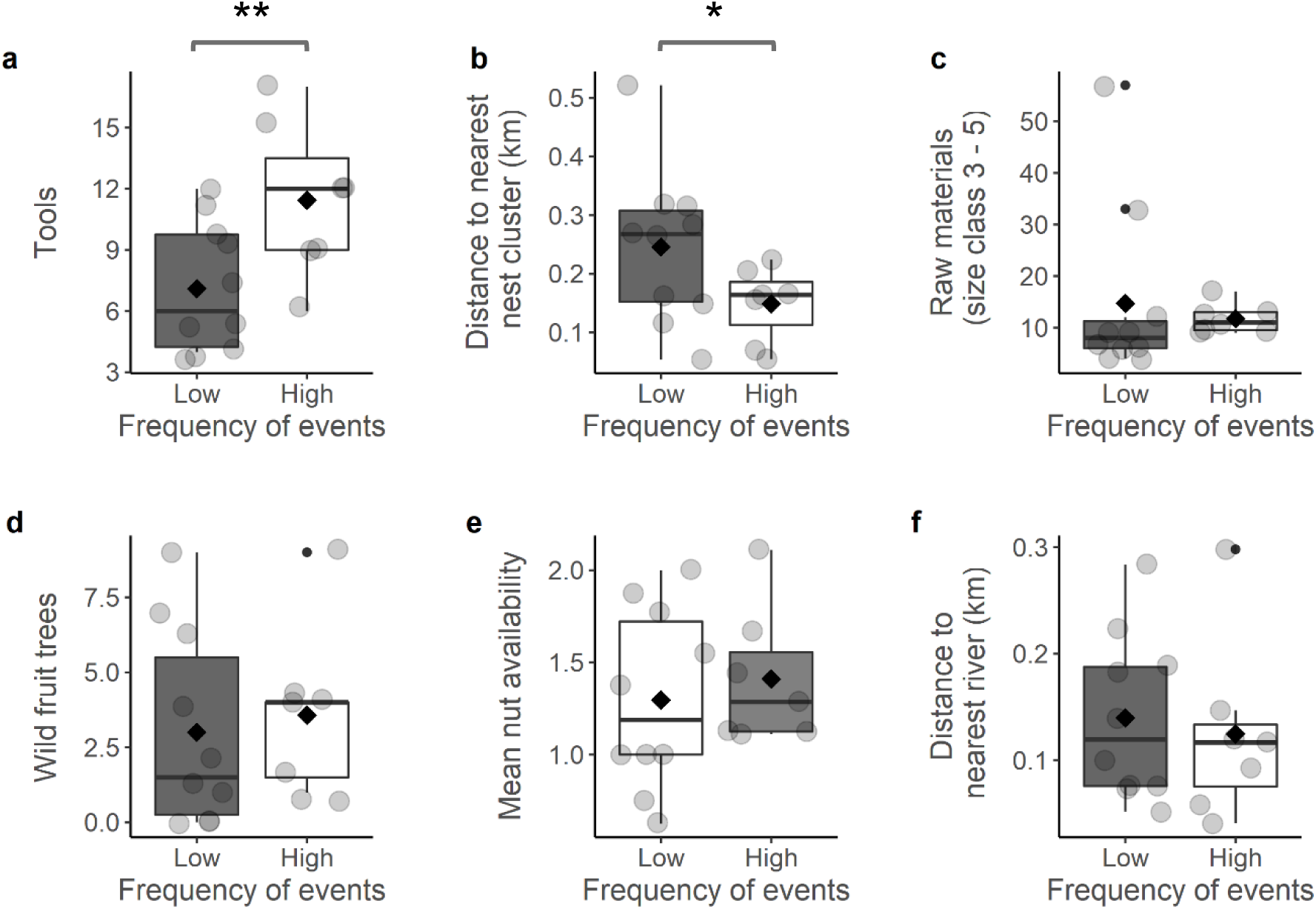
Frequency of tool site use in relative to: a) Raw materials that have been used as tools; b) Distance to nearest nest cluster (km); c) Raw materials of size class 3 to 5; d) Wild trees that are sourced by chimpanzees for food; e) Wild trees that are sourced by chimpanzees for fruit; f) Distance to nearest river (km). Grey circles represent individual points, and means indicated by diamonds. ** p < 0.05; * p < 0.07.

### Tool site inactivity

Out of the sub-models, the tool subset model yielded the best fit, although the AICc for the tool and fruit trees model was only marginally higher and produced comparable results (Appendix, Table A7, Table A8). Comparison of the tool subset model with the null model was significant (LRT: *df* = 3, *deviance* = 13.20, *p* < 0.01). Overall, we found that lower values of mean nut availability and a lower number of tools were both significant predictors of tool site inactivity, while wild food trees had no effect (Figure 6; Table 3). However, the data distributions shown in Figure 6 suggest that the model is not very robust.

**Figure 6.**
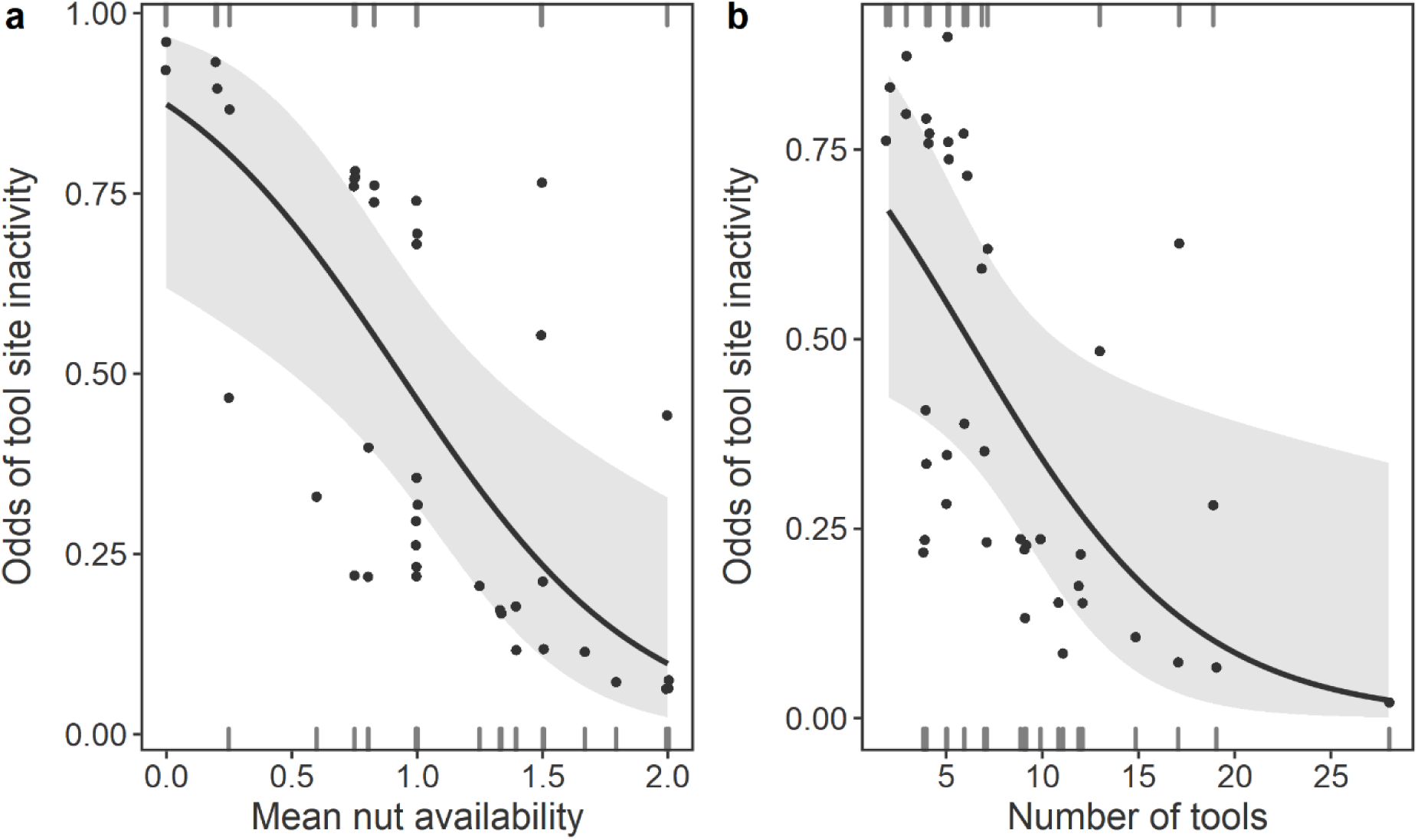
Probability of tool site inactivity in response to: a) Mean nut availability; b) Raw materials that have been used as tools.

**Table 3.** Results of the final GLM model investigating potential predictors influencing tool site abandonment

| Term | Estimate | SE | $\chi^2$ | p<sup>a</sup> |
| --- | --- | --- | --- | --- |
| Intercept | 2.957 | 1.263 |  | NI<sup>b</sup> |
| Mean. nut availability | -2.186 | 0.851 | 8.947 | 0.003 |
| Tools | -0.186 | 0.093 | 5.183 | 0.023 |
| Wild food trees | 0.093 | 0.109 | 0.752 | 0.386 |

<sup>a</sup> Results from the likelihood ratio test using the 'drop1' function.
<sup>b</sup> Not indicated because it has a limited interpretation.

## Discussion

### Tool site selection

From the initial inspection of the data, it is evident that a minimum of one oil palm, specifically an oil palm in close proximity (within 10 metres), is required for nut-cracking to occur in a given location. Further to this, our results show that the abundance of raw materials and food trees as well as proximity to the nearest nest cluster are also important predictors for whether a tool site is established at an oil palm location. This suggests that, in addition to the *ecological pre-requisites* of nut-cracking, i.e., a producing oil palm and raw materials for tools, other predictable resources that form part of the chimpanzee diet (wild food-providing trees), as well as non-food related activities (sleep sites), are influential in the spatial distribution of nut-cracking locations. In contrast, THV had a negative but non-significant effect on whether tool-sites occurred. While THV is a frequently consumed food item by the chimpanzees of Bossou (Humle, 2011a), it differs from other wild plant foods in that they have low calorific value and an individual plant often can only be sourced once, after which it is permanently depleted. In contrast, food trees are replenishable, often seasonally, and constitute reliable and predictable resources that can be returned to on a seasonal basis. The fact that THV does not bear a clear effect on tool site occurrence suggests that unpredictable food sources are not ecological drivers of tool site selection.

Previous research has found that many primate species have goal-orientated foraging trajectories towards spatially permanent resources and that they likely use mental maps to guide their resource exploitation strategies (Trapanese et al., 2019). Our results provide tentative evidence that the chimpanzees of Bossou may behave in a similar way, whereby nut-cracking activities take place within a foraging strategy that primarily targets predictable, high-value foods, while low-energy unpredictable foods like THV act as part of an opportunistic strategy during forage-on-the-go.

Distance to nearest nest cluster was a significant predictor in all models, whereby the likelihood of a tool site occurring increased with proximity to nest locations. Previous research has found that nests sites occur in areas of high food availability (Basabose and Yamagiwa, 2002; J. S. Carvalho et al., 2015; Furuichi and Hashimoto, 2004; Goodall, 1962; Janmaat et al., 2014). Given that nut-cracking sites are also located in areas with a greater number of food providing trees, and bearing in mind that the Bossou chimpanzees source oil palms for a range or other resources (e.g. fruit, pith, palm heart) and also nest in their crowns (Humle and Matsuzawa, 2001, 2004; Yamakoshi and Sugiyama, 1995), it is possible that the relationship between nut-cracking sites, proximity to nest sites, and food availability, is indicative that these areas are activity hotspots, rich in resources and with habitat characteristics that are suitable for a range of core chimpanzee activities.

Distance to the nearest river was not a significant predictor in any of the models. This contradicts previous research in the nearby forest of Diecké, that identified that nut-cracking locations occurred near waterlines (S. Carvalho et al., 2007). Emerging research on the role and importance of water in shaping primate behaviour, adaptations, and landscape use, is providing increasing evidence that there are differences in water-dependence between populations. Rainforest-dwelling apes can usually obtain their daily hydration requirements from the food they consume, and can go several days without drinking (Pontzer et al., 2021). However, for primates that live in year-round or seasonally arid landscapes water is a critical resource that shapes movement patterns and landscape use (Barton et al., 1992; Pruetz and Herzog, 2017; Wessling et al., 2018). Fongoli chimpanzees usually drink water at least once a day and often spend time near water sources during dry months to stay cool (Wessling, pers. comm.; Pruetz and Bertolani, 2009). Conversely, the Bossou forest is much more humid, with a long-wet season. It’s many streams, and small forest area provide a hydrological landscape in which chimpanzees are rarely more than 300 metres away from water. Furthermore, during a total of ∼500 hours of focal follows, the Bossou chimpanzees were only seen to drink water on eight occasions, suggesting that they can get most of their fluids from the foods they consume, in line with the general trend for non-human apes (Pontzer et al., 2021).

While water was not a significant factor for Bossou, we predict that it could be a major ecological driver regarding the spatial distribution and reuse of tool sites by savannah-living chimpanzees. The Fongoli chimpanzees do not crack nuts, but they engage in termite-fishing, which is also a spatially discrete technological activity tethered to the location of termite mounds (Bogart and Pruetz, 2008, 2011), much like nut-cracking.

Nevertheless, climatic or hydrological differences cannot explain the differences between Bossou and Diecké. Given the proximity of both field sites (approx. 50 km) and similar climates it is unlikely that this is due to differences in aridity or water availability. However, the chimpanzees of Diecké crack different nut species, *Panda oleosa* and *Coula edulis*, which are absent in Bossou and may be more water dependent than the oil palm. Thus, this discrepancy may be connected to the different plant species exploited and their respective ecology and distribution.

### Tool site use

Number of tools was the only variable of interest that differed significantly between sites with low or high frequency of nut-cracking events. This could indicate that the visible traces of nut-cracking found on tools act as visual cues for stimulating further nut-cracking behaviour. The repeated use of discrete locations through stigmergy has been suggested to have led the emergence of persistent places during the Middle Pleistocene (Matthew Pope et al., 2006; Matthew Pope, 2017; Shaw et al., 2016). Similar hypotheses featuring local and stimulus enhancement in chimpanzees have also been discussed as processes of social learning (e.g. in the development of technical skills; Musgrave et al., 2020; Tennie et al., 2020; Whiten, 2021) as well why some plants are sourced more intensively than others for the manufacture of termite fishing tools (Almeida-Warren et al., 2017). Conversely, it could indicate that chimpanzees prefer sites with material that they are already familiar with. Previous research in Bossou has demonstrated that chimpanzees reuse hammer-anvil pairs (tool-sets) more often than others, that there is both group- and individual-level preference for certain tool-sets (S. Carvalho et al., 2009), and that chimpanzees are selective of the types of materials they use for nut-cracking (S. Carvalho et al., 2008). Analogous studies on chimpanzee plant technologies, suggests similar patterns in the selection of materials for termite-fishing, ant-dipping, honey gathering and water extraction (Almeida-Warren et al., 2017; Lamon et al., 2018; Pascual-Garrido et al., 2012; Pascual-Garrido and Almeida-Warren, 2021).

Distance to nearest nest cluster showed a weak yet noteworthy difference, whereby the frequency of nut-cracking events was marginally greater at tool sites that were closer to nest locations. These results mirror those found for tool-site selection and offer further tentative support that active tool sites and their frequency of use is influenced by their distribution relative to current activity hotspots.

The number of wild food and fruit trees was largely the same for all active nut-cracking sites. This suggest that, while food providing trees are good indicators of tool-site selection, they may not good predictors of site use because the data collected did not capture temporal changes in food availability or frequency of foraging activity. On the other hand, nests are temporary features that rarely preserve for longer than six months in non-savannah environments (Ihobe, 2005; Kamgang et al., 2020; Zamma and Makelele, 2012). Therefore, they are a better spatial proxy for recent ranging patterns and possibly explains why differences were found for nests, but not for vegetation.

Some consideration needs to be given as to the low number of weekly traces of nut-cracking events recorded per tool site during the 15-weeks of monitoring. This is partially due to the fact that not all active tool sites were monitored, with a further seven traces were found through indirect observations at non-monitored sites. Our data indicates that a minimum of 40 nut-cracking events took place at natural nut-cracking sites during the 15 week monitoring period, averaging approximately three events per week, which may be sufficient for the existing chimpanzee population.

Furthermore, out of the nine nut-cracking events witnessed during group follows, six took place at the outdoor laboratory, where nuts and stones were being artificially provisioned for another project. The outdoor laboratory is located at the intersection of several routes which the chimpanzees frequently travel through to access different parts of the forest. As a location that has always experienced a high degree of natural thoroughfare (Tetsuro Matsuzawa, 2011), it may represent a pre-existing activity hotspot that has been enhanced by the guaranteed encounter of tools and edible nuts. This could explain why a comparatively higher number of nut-cracking events were observed there, similar to patterns recorded by Hockings et al. (2009).

Nevertheless, due to small sample sizes and the limitations of the statistics employed, more data are needed to explore these hypotheses further. For future research, it would be important to investigate over a longer timescale whether and how often chimpanzees visited the part of the forest where the tool site is located. This could make the use of complementary data from camera traps placed in strategic locations, as full-day focal follows are not permitted in Bossou.

### Tool site inactivity

Understanding the contexts of tool site inactivity is an important step in investigating the conditions required for nut-cracking to occur and persist over time in a particular location, and the factors that might lead to their abandonment. Our data suggests that mean nut availability, used as a proxy for tree productivity, and a high abundance of tools are important in maintaining the active status of a nut-cracking site. However, there are clear exceptions that appear to not quite fit the model (Figure 6), suggesting that other factors that were not considered in the analysis may also be at play.

The Bossou forest suffers from a great deal of human activity, particularly slash-and-burn agriculture, which leads to frequent and rapid changes in the spatial distribution of resources and localized vegetation composition (Hockings, 2011). While oil palms are not cut down during this process and are highly resistant to fire (Yamakoshi, 2011), the changes in the surrounding landscape and the increase in human presence may deter chimpanzees from visiting those areas, especially if they are near the forest boundary. Conversely, cultivated land that contains desirable food items (e.g. banana, mango, papaya) can often attract chimpanzees (Hockings, 2011), and perhaps, under these conditions, the chimpanzees prioritize the prized fruit over nuts that can be found almost anywhere. Site inactivity could also be an artefact of population decline, whereby fewer resources are sufficient to sustain the entire population. Previous literature has suggested that the Bossou forest has a carrying capacity for around 20 chimpanzees (Sugiyama and Fujita, 2011), so it is possible that the current population may no longer need to depend as highly on nuts to supplement their diets. A future longitudinal comparison drawing from historical and contemporary data will help investigate and test this further.

## Conclusions

Our results indicate that proximity to a nut tree, an abundance of raw materials and predictable resources, as well as proximity to a nesting site are important ecological parameters for the establishment of a nut-cracking site in a given location. Distance to nearest nest cluster was also correlated with frequency of nut-cracking, which could potentially indicate that nesting sites are important anchors for ranging and activity patterns. Similarly, tool availability was significantly correlated with tool site use, as well as tool site inactivity, suggests that familiarity of materials used for tools or the visual cues of tool use could be important in the persistence of nut-cracking activities once a site has been established. While there was no significant difference in nut availability among oil palms at active sites, the odds of tool site inactivity were greater when mean nut availability was low, potentially indicating that a decline in oil palm productivity at nut-cracking sites is driver of sites disuse. Together, these results postulate that nut-cracking in Bossou is not only tethered to locations that provide the necessary resources for this activity but is also intimately connected to a broader foraging and behavioural landscape that is mediated by the spatio-temporal availability of primary target resources, such as predictable food-providing trees, as well as the distribution of frequently used nesting locations.

Preliminary comparisons with other sites regarding the importance of ecological features such as the effect of water on tool use and ranging patterns, suggests that the ecology of chimpanzee technology is context-specific and should be examined with this in mind. Further studies investigating the technological landscapes of other chimpanzee populations, as well as the integration of long-term data, will help better understand the effect of different environmental and demographic contexts on the factors driving the spatial distribution and reuse of tool sites, adding further detail to this picture.

While current evidence suggests that early hominin and chimpanzee lithic technologies differ in form and function (Arroyo and de la Torre, 2016; Toth et al., 2006), it is likely they had similar plant-dominated diets supported by insects and sporadic meat consumption (Panger et al., 2003). Thus, it is plausible that, like chimpanzees, early hominin tool-use operated within behavioural landscapes conditioned by localized environmental parameters, where foraging strategies were shaped by the distribution and availability of predictable food sources, the dietary dependence on extractive foraging and the availability of the necessary raw materials, as well as the location of safe places for sleeping. With the aid of primate archaeological inference, visualizing the spatial distribution of hominin lithic assemblages within this framework will be instrumental in providing crucial insights for reconstructing the patterns of landscape use and resource exploitation.

The present study suggests that the technological landscape of the chimpanzees of Bossou shares affinities with the ‘favoured places’ model (Schick and Toth, 1993; Shick, 1987), which proposed that hominin tool sites formed at the centre of foraging areas where hominins would process and consume food, rest and socialize, with sites being used more intensively in areas with higher resource abundance. Such ‘activity hotspots’ would have acted as ecological tethers, shaping early hominin movement and foraging patterns, which, in turn, would have led to the formation and repeated use of tool sites over time. As the most conspicuous evidence of these locations, stone tool assemblages may hold important clues for uncovering behaviours beyond those associated to lithic technology and serve as starting points to search for traces of other activities, such as sleeping, foraging, and insectivory, that are currently extremely rare in the archaeological record. This research draws upon the work of Glynn Isaac (e.g. Isaac, 1981; Isaac et al., 1981; Isaac and Harris, 1980) who pioneered the application of landscape-scale approaches to the study of hominin assemblages, from which the first concrete models of hominin site formation were developed. Further studies will help guide future human origins research and provide an empirical framework for modelling and testing hypotheses of early hominin behaviour associated to the archaeological record of our earliest ancestors.

## Acknowledgements

We are grateful to Vincent Mamy, Lawe Goigbe, Boniface Zogbila, Gouanou Zogbila, Jules Doré, Henry Camara, and Pascal Goumi for support in the field; Dr. Jesse Van der Grient and Dr. Alexander Mielke for statistical advice; Dr. Dora Biro and Dr. Bronwyn Tarr for helpful comments and feedback on the original manuscript. We also thank the Direction Nationale de la Recherche Scientifique, the Institut de Recherche Environmentale de Bossou (Guinea), and the Kyoto University Primate Research Institute, for research permissions and logistical support. KAW was funded by the Fundação pela Ciência e Tecnologia (grant number: SFRH/BD/115085/2016), supported by the Programa Operacional Capital Humano (POCH) and the European Union; the Boise Trust Fund (University of Oxford); and the National Geographic Society (grant number: EC-399R-18); SC thanks the Leverhulme Trust for the PLP 2016-114 which supported part of the field equipment and logistics; TM thanks the Japan Society for the Promotion of Science (JSPS) Leading Graduate Program (U04-PWS), JSPS core-to-core CCSN, and the Ministry of Education, Culture, Sports, Science and Technology of Japan (MEXT)/JSPS-KAKENHI (grant numbers: #07102010, #12002009, #16002001, #20002001, #24000001, #16H06283).

## Author contributions

KAW is the main author and contributor and was responsible for conceptualization, methodology, data collection, formal analysis, data curation, visualization and writing of the original manuscript. TM provided resources and editorial advice. SC provided supervision, data and resources and contributed towards conceptualization, methodology and revisions of the manuscript.

## Appendix

**Table A1.** Summary of the response variables used in this study and their descriptions.

| Variable | Description | Units | Range |
| --- | --- | --- | --- |
| Raw materials | Total number of stones in the survey quadrat | Counts | 0-99 |
| Raw materials<br>(size class 3-5) | Number of stones with sizes within the 95% confidence interval of tools used of nut-cracking | Counts | 0-81 |
| Tools | Total number of stones used for cracking nuts; Excludes unused fragments | Counts | 0-28 |
| Food trees | Total number of plants know to provide food sources to Bossou chimpanzees; Excludes plants with diameter at breast high (DBH) under 2 cm | Counts | 0-12 |
| Fruit trees | Total number of plants know to be sources of fruit to Bossou chimpanzees; Excludes plants with diameter at breast high (DBH) under 2 cm | Counts | 0-9 |
| THV | Total number of individual terrestrial herbaceous vegetation (THV) plants that are part of the Bossou chimpanzee diet | Counts | 0-10 |
| Nut availability | Approximate amount of nuts with edible kernels; Mean nut availability used in analysis | Categorical | 0 (none); 1 (< 50); 2 (50 – 100); 3 (> 100) |
| Distance to nearest nest cluster | Distance, in metres, to nearest cluster of 21 nests; Cluster defined as nests within a 50-metre radius | Metres | 50.01-781.06 |
| Distance to nearest river | Distance, in metres, to nearest water course | Metres | 6.92-399.84 |

**Table A2.** Likelihood ratio test results of the Full-Null model comparisons of the site selection models.

| Model | Resid. df | Resid. Dev | df | Deviance | p |
| --- | --- | --- | --- | --- | --- |
| ~1 (Null model) | 81 | 113.627 |  |  |  |
| ~Raw materials + Wild food trees + THV + Nest +River | 76 | 59.289 | 5 | 54.338 | <0.001 |
| ~**Raw materials (size class 3-5)** + Wild food trees + THV + Nest +River | 76 | 57.109 | 5 | 56.519 | <0.001 |
| ~Raw materials + **Wild fruit trees** + THV + Nest +River | 76 | 62.480 | 5 | 51.147 | <0.001 |
| ~Raw materials (size class 3-5) + Wild fruit trees + THV + Nest +River | 76 | 61.817 | 5 | 51.810 | <0.001 |

**Table A3.**
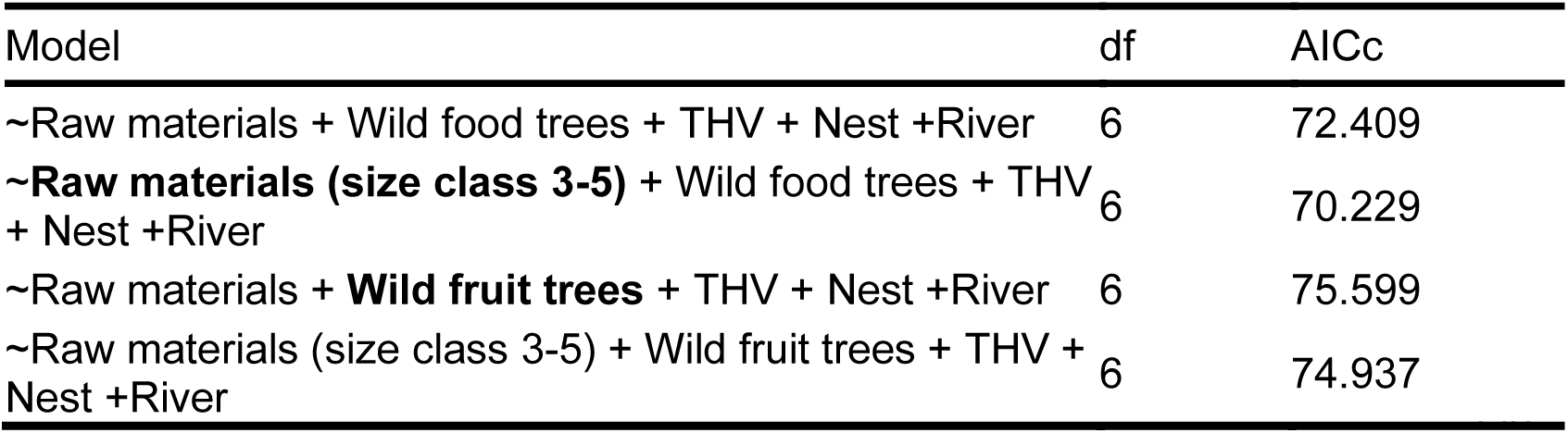
Results of the AICc-Based Model Selection of the GLM for the predictors of tool site selection. Subset variables highlighted in bold.

| Model | df | AICc |
| --- | --- | --- |
| ~Raw materials + Wild food trees + THV + Nest +River | 6 | 72.409 |
| ~**Raw materials (size class 3-5)** + Wild food trees + THV + Nest +River | 6 | 70.229 |
| ~Raw materials + **Wild fruit trees** + THV + Nest +River | 6 | 75.599 |
| ~Raw materials (size class 3-5) + Wild fruit trees + THV + Nest +River | 6 | 74.937 |

**Table A4.** Results of the additional GLMs investigating potential predictors of tool site selection. Subset variables highlighted in bold.

| Model | Term | Estimate | SE | $\chi^2$ | P<sup>a</sup> |
| --- | --- | --- | --- | --- | --- |
| ~Raw materials + Wild food trees + THV + Nest + River | Intercept | -0.575 | 1.076 |  | NI<sup>b</sup> |
|  | Raw materials | 0.195 | 0.057 | 19.748 | <0.001 |
|  | Wild food trees | 0.303 | 0.155 | 4.944 | 0.026 |
|  | THV | -0.101 | 0.159 | 0.419 | 0.518 |
|  | Distance to nearest nest cluster | -0.007 | 0.003 | 7.377 | 0.007 |
|  | Distance to nearest river | 0.004 | 0.004 | 0.887 | 0.346 |
| ~Raw materials + **Wild fruit trees** + THV + Nest + River | Intercept | -0.305 | 1.052 |  | NI<sup>b</sup> |
|  | Raw materials | 0.203 | 0.059 | 20.316 | <0.001 |
|  | Wild fruit trees | 0.241 | 0.193 | 1.753 | 0.185 |
|  | THV | -0.031 | 0.136 | 0.051 | 0.821 |
|  | Distance to nearest nest cluster | -0.008 | 0.003 | 8.469 | 0.004 |
|  |  | 0.005 | 0.004 | 1.270 | 0.260 |
| ~Raw materials (size class 3-5) + **Wild fruit trees** + THV + Nest + River | Intercept | -0.700 | 1.079 |  |  |
|  | Raw materials (size class 3-5) | 0.351 | 0.102 | 20.978 | 0.000 |
|  | Wild fruit trees | 0.199 | 0.202 | 1.049 | 0.306 |
|  | THV | -0.017 | 0.141 | 0.014 | 0.905 |
|  | Distance to nearest nest cluster | -0.007 | 0.003 | 7.715 | 0.005 |
|  | Distance to nearest river | 0.006 | 0.004 | 1.967 | 0.161 |

<sup>a</sup> Results from the likelihood ratio test using the 'drop1' function.
<sup>b</sup> Not indicated because it has a limited interpretation.

**Table A5.** Descriptive statistics and test results of the comparison between tool sites with low frequency (N = 10) and high frequency (N = 7) of nut-cracking events.

| Variable | Low frequency |  | High frequency |  | Test results |  |  |  |
| --- | --- | --- | --- | --- | --- | --- | --- | --- |
|  | Mean | SD | Mean | SD | Test<sup>a</sup> | t | df | p |
| Tools | 7.100 | 3.143 | 11.429 | 3.780 | tt | 2.487 | 11.429 | 0.029 |
| Distance to nearest nest cluster (m) | 245.660 | 132.760 | 148.714 | 64.047 | tt | -2.000 | 13.707 | 0.066 |
| Raw materials (size class 3-5) | 8.000<sup>b</sup> | 5.250<sup>c</sup> | 11.000<sup>b</sup> | 3.500<sup>c</sup> | wt | 50.000<sup>d</sup> | -1.430<sup>e</sup> | 0.154 |
| Raw materials | 8.000<sup>b</sup> | 14.500<sup>c</sup> | 14.000<sup>b</sup> | 4.000<sup>c</sup> | wt | 49.000<sup>d</sup> | -1.320<sup>e</sup> | 0.186 |
| Mean nut availability | 1.296 | 0.489 | 1.410 | 0.371 | tt | 0.547 | 14.835 | 0.593 |
| Wild fruit trees | 3.000 | 3.300 | 3.571 | 2.760 | tt | 0.387 | 14.401 | 0.704 |
| Distance to nearest river (m) | 139.613 | 77.486 | 124.710 | 84.836 | tt | -0.369 | 12.265 | 0.718 |
| Mean fruit bunches | 1.252 | 0.630 | 1.188 | 0.630 | tt | -0.204 | 13.069 | 0.842 |
| Wild food trees | 4.000 | 3.916 | 4.286 | 2.690 | tt | 0.178 | 15.000 | 0.861 |

<sup>a</sup> Test used: tt =t-test; wt = Wilcoxon rank sum test
<sup>d</sup> Reported as median
<sup>e</sup> Reported as inter-quartile range (IQR)
<sup>b</sup> Reported as w-statistic for Wilcoxon rank sum tests
<sup>c</sup> Reported as z-score for the w-statistic

**Table A6.** Likelihood ratio test results of the Full-Null model comparisons of the site inactivity models.

| Model | Resid. df | Resid. Dev | df | Deviance | p |
| --- | --- | --- | --- | --- | --- |
| ~1 (Null model) | 81 | 113.627 |  |  |  |
| ~Mean nut availability + Raw materials + Wild food trees | 36 | 46.532 | 3 | 8.017 | 0.046 |
| ~Mean nut availability + **Raw materials (size class 3-5)** + Wild food trees | 36 | 46.518 | 3 | 8.030 | 0.045 |
| ~Mean nut availability + **Tools** + Wild food trees | 36 | 41.350 | 3 | 13.198 | 0.004 |
| ~Mean nut availability + **Raw materials (size class 3-5)** + **Wild fruit trees** | 36 | 46.621 | 3 | 7.928 | 0.048 |
| ~Mean nut availability + **Tools** + **Wild fruit trees** | 36 | 41.968 | 3 | 12.581 | 0.006 |

**Table A7.**
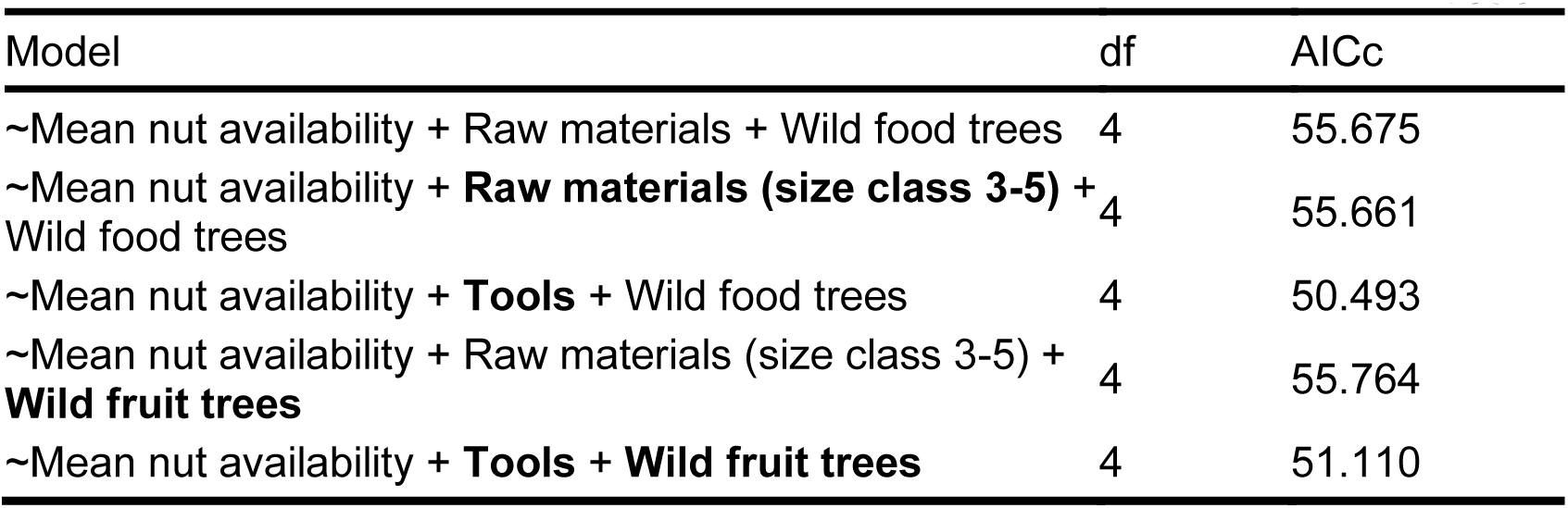
Results of AICc-Based Model Selection of the GLM for the predictors of tool site inactivity. Subset variables highlighted in bold.

**Table A8.** Results of the additional GLM models investigating potential predictors of tool site inactivity. Subset variables highlighted in bold.

| Model | Term | Estimate | SE | $\chi^2$ | P<sup>a</sup> |
| --- | --- | --- | --- | --- | --- |
| ~Mean nut availability +<br>Raw materials + Wild<br>food trees | Intercept | 1.275 | 0.927 |  |  |
|  | Mean nut availability | -1.909 | 0.788 | 7.757 | 0.005 |
|  | Raw materials (size<br>class 3-5) | 0.001 | 0.017 | 0.001 | 0.973 |
|  | Wild food trees | 0.083 | 0.111 | 0.567 | 0.452 |
| ~Mean nut availability +<br>**Raw materials (size**<br>**class 3-5)** + Wild food<br>trees | Intercept | 1.310 | 0.923 |  | NI<sup>b</sup> |
|  | Mean nut availability | -1.911 | 0.787 | 7.796 | 0.005 |
|  | Raw materials (size<br>class 3-5) | -0.003 | 0.022 | 0.015 | 0.903 |
|  | Wild food trees | 0.086 | 0.109 | 0.625 | 0.429 |
| ~Mean nut availability +<br>Raw materials (size class<br>3-5) + **Wild fruit trees** | Intercept | 1.838 | 0.971 |  | NI<sup>b</sup> |
|  | Mean nut availability | -1.895 | 0.773 | 7.757 | 0.005 |
|  | Raw materials (size<br>class 3-5) | 0.004 | 0.022 | 0.036 | 0.849 |
|  | Wild fruit trees | -0.104 | 0.145 | 0.522 | 0.470 |
| ~Mean nut availability +<br>**Tools + Wild fruit trees** | Intercept | 3.282 | 1.268 |  | NI<sup>b</sup> |
|  | Mean nut availability | -2.077 | 0.812 | 8.491 | 0.004 |
|  | Tools | -0.170 | 0.088 | 4.689 | 0.030 |
|  | Wild fruit trees | -0.053 | 0.144 | 0.135 | 0.714 |

<sup>a</sup> Results from the likelihood ratio test using the 'drop1' function.
<sup>b</sup> Not indicated because it has a limited interpretation.

